# Palatability of Insecticides and Protein in Sugar Solutions to Argentine Ants

**DOI:** 10.1101/2024.08.29.610285

**Authors:** Thomas Wagner, Moana Vorjans, Elias Garsi, Cosmina Werneke, Tomer J. Czaczkes

## Abstract

Invasive ant species like *Linepithema humile* cause significant ecological and economic harm, making effective control strategies essential. Insecticide baits are currently the most effective approach for controlling ants. Therefore, quantifying how palatable or unpalatable baits, bait additives, or toxicants are, is critical for developing effective control methods. It has recently been demonstrated that in the comparative evaluation of foods, animals that are aware of both a test food and a comparator food exhibit greatly increased sensitivity when detecting the unpalatability of liquid baits. Here, we deploy a newly developed comparative evaluation methodology to examine the palatability to *L. humile* workers of three toxicants used in invasive ant control: Fipronil, spinosad, and imidacloprid, as well as egg white protein.

Ants showed no significant preference between pure sucrose and sucrose-toxicant solutions, indicating that they either cannot detect the toxicants or that they do not find them distasteful. Survival tests confirmed that the toxicant concentrations used were lethal, with a survival rate of 50% or below after 72 hours. However, ants found egg protein additive unpalatable, significantly preferring pure sucrose to a sucrose-egg protein mix.

These findings confirm that three major toxicants are suitable for use in baits, and that reported abandonment or avoidance of toxic baits is not due to the unpalatability of these toxicants. However, the addition of egg protein to sucrose baits, even at ratios which optimise colony growth, is likely counterproductive. Future research should investigate the relative preference of invasive ants for various bait matrixes over naturally available food, ensuring more effective pest management strategies.

## Introduction

Invasive ant species cause significant economic harm and severe ecological disruption (Gutrich et al. 2007; Siddiqui et al. 2021; Angulo et al. 2022). Controlling invasive ant species has thus become a top priority for pest management professionals worldwide. Unfortunately, combating invasive ants has often proven to be challenging. For example, efforts to control the Argentine ant (*Linepithema humile*) or the red imported fire ant (*Solenopsis invicta*) have frequently failed, with populations rebounding after treatment (Silverman and Brightwell 2008; Hoffmann et al. 2016; Buczkowski 2024).

One promising approach involves using baits with slow-acting toxicants, which allow time for the toxicant to reach the queen via the workers. Studies have shown that hydrogels or granular baits combined with slow-acting insecticides can effectively deliver toxicants to the queen, resulting in more comprehensive colony control (Hoffmann 2011; Buczkowski et al. 2014; Rust et al. 2015; Sunamura et al. 2022). However, baits are not always consistently effective. For example, a study using fipronil-based baits for red imported fire ant control found that although initial reductions in ant populations were observed, the ants quickly rebounded, indicating that the bait did not provide long-term control (Ipser and Gardner 2010).

Moreover, baits can be rejected over time as ants often prefer natural high-quality food sources, which act as competitive food sources and divert foraging away from the baits (Rust et al. 2003; Silverman and Brightwell 2008). Ants have also been shown to actively abandon harmful food (Herz et al. 2008; Arenas and Roces 2016a; Zanola et al. 2024). Understanding the feeding preferences of ants is crucial for designing baits that are attractive to and consumed by ants, especially when they contain various ingredients such as toxicants, bittering agents, co-formulants, and attractants. If the bait and its components do not align with the ants’ preferences, it may be rejected or less preferred compared to other available food sources. Therefore, it is essential to understand what ants like or dislike when developing more effective pest control strategies.

To investigate these preferences, we utilized a recently developed dual-feeder method (Wagner et al. 2024). This technique allows for the simultaneous presentation of two bait formulations, allowing ants to directly compare the two options. Comparative evaluation is a more sensitive approach to quantifying ant feeding preference than absolute food acceptance or cafeteria-style approaches, as indicated in Wagner et al (2024). Additionally, the dual feeder method avoids inducing neophobia or the effects of sequential exposure, both of which have been documented in ants (Oberhauser and Czaczkes 2018; Oberhauser et al. 2022).

This method is particularly relevant because, although ant feeding preferences in nature are probably mostly initially sequential, they become comparative over time as ants gain experience with multiple food sources. Most relevant for pest control is the comparison ants will make between natural food sources and the bait. Thus, understanding comparative preferences provides a more realistic, as well as more sensitive, assessment of bait efficacy than absolute food acceptance tests. Ants are highly responsive to their environment and adjust their decisions of what food to collect through colony feedback or individual and collective comparative evaluation (Detrain and Deneubourg 2006; Dussutour and Simpson 2008; Arganda et al. 2014; Price et al. 2016).

Bait evaluations often focus on whether ants consume the bait without initially considering whether the ants can detect the toxicant but still consume the bait due to its initial palatability or a lack of alternative options (Silverman and Roulston 2001; O’Brien 2005; Lach et al. 2019). Understanding whether ants can perceive the insecticides may help explain why baits might be abandoned over time, as ants may initially accept the bait but avoid it later if they detect the insecticide and have access to alternative food sources.

The primary aim of our study was to assess whether Argentine ants could detect and thus avoid sucrose solutions mixed with three different insecticides—fipronil, spinosad, and imidacloprid—by evaluating their acceptance and consumption of these treated solutions compared to untreated sucrose. These insecticides are some of the most commonly used toxicants for invasive ant control (Rust et al. 2003; Klotz et al. 2007; Hoffmann et al. 2016; Buczkowski and Wossler 2019). Fipronil is a broad-spectrum insecticide that disrupts the central nervous system of insects. Spinosad also affects the nervous system of insects but has a lower toxicity to non-target organisms, especially fish. Imidacloprid also affects the nervous systems of insects and is widely used in agriculture against a variety of insect pests. All three insecticides have been reported to successfully eradicate ants locally in certain situations (Rust et al. 2003; Ipser and Gardner 2010; Khan et al. 2021; Subekti et al. 2022).

Secondly, we aimed to determine whether ants prefer pure sucrose solutions over sucrose solutions mixed with protein. While previous studies, such as Buczkowski and Krushelnycky (2012), have shown that a blend of protein and carbohydrate can be highly attractive to a variety of ant species, including Argentine ants, our study seeks to directly compare the preference for pure sucrose versus a protein-sucrose mix. This comparison is crucial because, although ants may find a protein-carbohydrate blend attractive, they might still prefer pure carbohydrate baits under certain conditions. Understanding these preferences can provide insights into the most effective bait formulations. For instance, adding protein to a sucrose bait might increase palatability for some ants, but if they exhibit a stronger preference for pure sucrose, it could suggest that pure carbohydrate baits might be more effective in certain scenarios. Furthermore, ants are known to forage according to specific nutrient intake ratios, seeking out proteins and carbohydrates as needed to achieve this balance (Arganda et al. 2014; Csata et al. 2020) Therefore, testing whether a pure carbohydrate bait is preferred over a protein-sucrose mix is essential to optimizing bait effectiveness

By analysing the feeding behaviour of invasive ants, we aimed to gain insights into their nutritional preferences and how these preferences could be leveraged to develop more effective baits.

## Materials & Methods

### Colony Fragments Maintenance

*Linepithema humile* were sourced from Girona, Spain (February 2023), and Proença-a-Nova, Portugal (May 2022). These ants were all part of the same European supercolony, but they were collected from different locations within that region. Colony fragments were established, each comprising one or more queens, brood and ranging from 300 to 1000 workers. These colonies were housed in plastic foraging boxes (32.5 cm x 22.2 cm x 11.4 cm), with a plaster of Paris floor. The walls were coated in a Talc-EtOH mixture (20g Talc in 100ml EtOh 20%) to prevent escape (Ning et al. 2019). Within each box, multiple 15 ml plastic tubes were provided as nests, covered with red transparent plastic foil, partially filled with water, and sealed with cotton plugs.

The colonies were maintained under a 12:12 light:dark cycle at room temperature (24-26°C), with continuous access to water via both the sealed tubes and a water feeder. Additionally, colonies were supplied ad *libitum* with a 0.5M sucrose solution and freeze-killed *Drosophila melanogaster*, with food withheld for 4 days prior testing.

#### Solution and Insecticide composition

We prepared a 1M sucrose-water solution (Südzucker AG, Mannheim, Germany) and 1M sucrose-water solutions with varying concentrations of acetone (purity 99.5%, Sigma-Aldrich, Taufkirchen, Germany) at 0.052%, 0.12%, 0.52%, and 1.2% v/v. These solutions served as counterparts for the toxicant solutions, as some of the toxicants are soluble in acetone but not water. Three toxicants were used: imidacloprid (purity 99.7%, CAS 138261-41-3, Sigma-Aldrich, Taufkirchen, Germany), fipronil (purity 99.7%, CAS 120068-37-3, Sigma-Aldrich, Taufkirchen, Germany), and sinosad (purity 99.4%, CAS 168316-95-8, Sigma-Aldrich, Taufkirchen, Germany).

The concentrations of the insecticides were selected based on their typical application levels in commercial baits: fipronil at 0.0001% and 0.001% (Nyamukondiwa and Addison 2011, 2014), spinosad at 0.015% and 0.15% (Daane et al. 2008), and imidacloprid at 0.005% (Rust et al. 2003).

Imidacloprid has good water solubility and was thus mixed with a 1M sucrose solution to create a 1M + imidacloprid 0.005% solution. However, spinosad and fipronil have poor water solubility, so acetone was added to improve solubility. We mixed spinosad with acetone to create a 1M + spinosad 0.15% and 1.2% acetone solution, a 1M + spinosad 0.015% and 0.12% acetone solution, and a 1M + fipronil 0.001% and 0.52% acetone solution, as well as a 1M + fipronil 0.0001% and 0.052% acetone solution.

Additionally, a 1M sucrose solution mixed with egg protein (egg white isolate from LeeGroup GmbH) in a 25:1 ratio of sucrose to protein was created. We chose this ratio based on preliminary laboratory tests, which indicated that *L. humile* highly accept and consumes it (H. Galante, unpublished data), and discussion with experts on *L. humile* nutrition [A. Dussutour & E. Csata, personal communications, see also Arganda et al. (2014)].

#### Experiment procedure –Assessing perception and preference in a dual-feeder system

The primary measure examined was the proportion of time spent drinking from the first-choice feeder solution. This metric was chosen because ants that fully accept a liquid food source typically continue feeding until they are satiated (Wendt and Czaczkes 2017; Wagner et al. 2024).

The experimental setup followed the design described in Wagner et al. (2024). A single ant was allowed to walk onto a piece of paper and then walk off it onto a 10 cm long runway, covered in replaceable paper overlays (to facilitate pheromone trail removal), leading to the dual-feeder landing platform (figure 1). The dual-feeder features two triangular wells converging into parallel channels, each 0.35 mm in width, which efficiently transport liquids to their ends via capillary action. A narrow gap of 0.30 mm between the channels ensures that an ant, upon first contact with one solution, will almost immediately encounter the alternate solution with their antennae, facilitating a rapid and informed choice between the two options. Comparative evaluation of food sources is more sensitive in ants than absolute evaluation (Wagner et al. 2024). The following solution pairs were tested:

- 1M sucrose vs. 1M sucrose + imidacloprid (0.005% concentration),
- 1M sucrose + acetone (0.52%) vs. 1M sucrose + acetone (0.52%) + fipronil (0.001%),
- 1M sucrose + acetone (0.052%) vs. 1M sucrose + acetone (0.052%) + fipronil (0.0001%),
- 1M sucrose + acetone (1.2%) vs. 1M sucrose + acetone (1.2%) + spinosad (0.15%),
- 1M sucrose + acetone (0.12%) vs. 1M sucrose + acetone (0.12%) + spinosad (0.015%).
- 1M sucrose vs. 1M sucrose + egg-white-protein (25:1 ratio)

**Figure 1:**
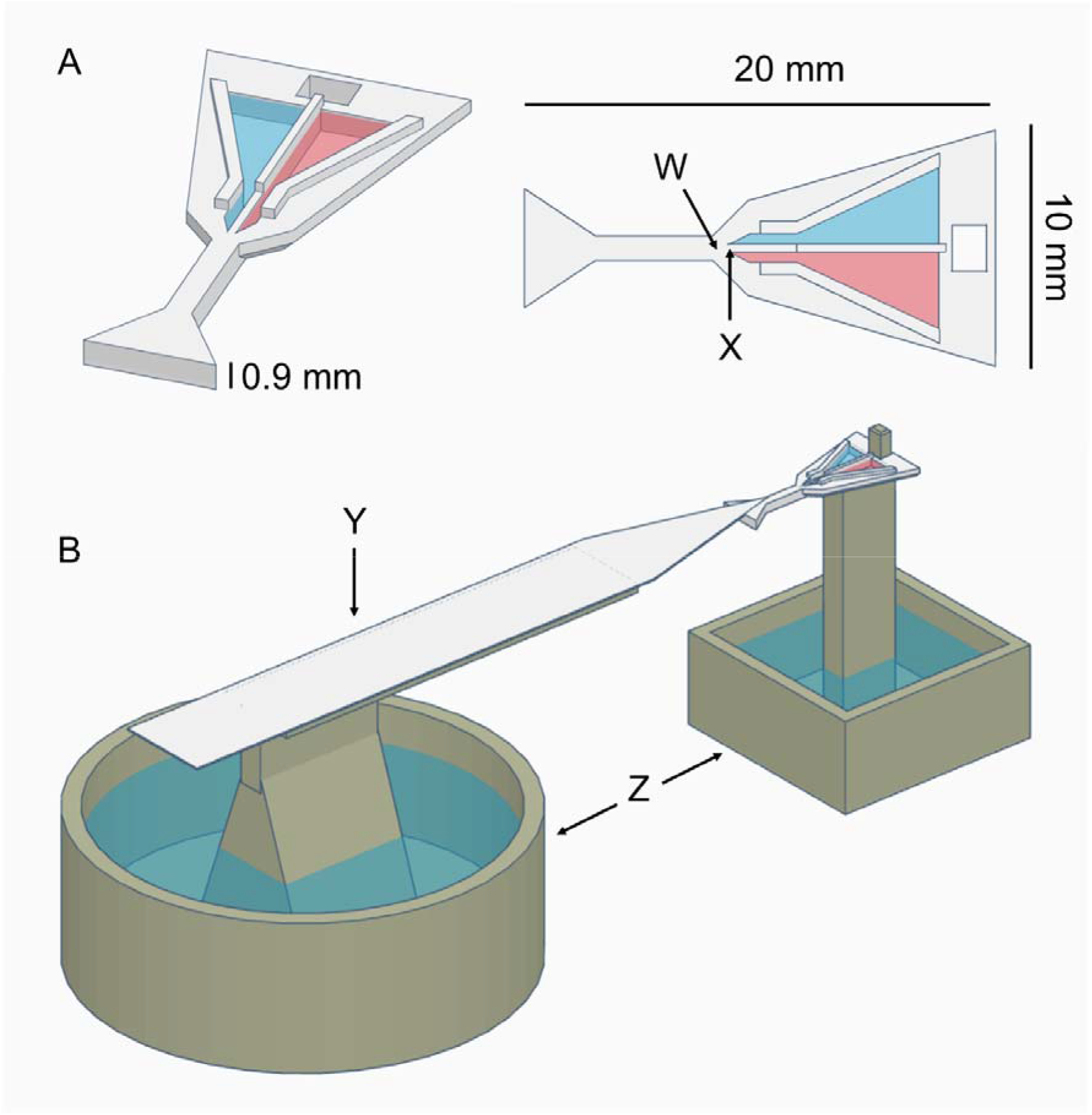
Dual-Feeder, following Wagner et al. (2024): A: Dual-feeder, W: Optimal drinking position, X: 0.30mm gap separation between the two wells containing solution 1 and 2 (here visualised as blue and red), B: Whole experimental setup showing the dual-feeder connected with the runway Y and attached to a pillar, surrounded by a water moat Z.

The side of the well containing the different solutions was alternated between ants. Ants initially encountered one of the drinking wells, either with their mandibles or antennae, and began drinking. They could easily and quickly switch to the other well by simply moving their head. The side initially encountered and the pure drinking time for each solution were recorded using a stopwatch, measuring only continuous drinking and excluding interruptions or side switches. The duration of each trial was capped at 5 minutes or until the ant returned to the runway, indicating it was satiated. A total of 498 ants from 16 distinct colony fragments were tested.

##### Assessment of Ant Survival Rate

The assessment of ant survival rate was conducted to demonstrate that the toxicant doses were sufficient to cause mortality over a 72-hour period, confirming their efficacy for pest control purposes in natural settings (Rust et al. 2003). However, it should be noted that in natural conditions, ants may consume poisoned bait multiple times, or unload the bait to their nestmates

All 498 ants tested above (from 16 colony fragments) were transferred to plastic petri dishes (Ø14 cm x 2 cm) immediately after testing. A centrifuge tube containing water, sealed with cotton, was provided in each dish over the 72 hours. Ants had no access to the baits after testing. The petri dishes were then placed in a temperature-controlled room maintained at 24-26°C, with a 12:12 light cycle. Ant survival was assessed at 24-hour intervals for 72 hours. Control petri dishes containing ants, previously fed with 1M sucrose, and a centrifuge tube containing water and sealed with cotton were included for comparison. The number of ants dead at each time interval was recorded. Survival was not recorded for the ants tested for protein preference.

In addition, to test for any mortality effect of the acetone used to increase toxicant solubility, groups of ants (6 groups of 16 ants) were allowed to feed to satiation on either pure 1M sucrose or 1M sucrose and 1.2% acetone. Survival was recorded as above.

##### Statistical analysis

The full statistical analysis results, as well as the complete dataset upon which this analysis is performed, can be accessed at https://zenodo.org/records/13482777. All statistical outputs were generated using R version 4.2.2 (R Core Team 2022), with figures produced using ggplot2 (Wickham 2016). Binomial generalized linear mixed-effects models (GLMM) (Bates et al. 2018) were used to test for differences in time spent feeding at various food sources relative to the control food. DHARMa (Hartig 2021) was used to assess linear model assumptions. Estimated marginal means and contrasts were derived using the emmeans package (Lenth et al. 2024) with Tukey corrections applied to account for multiple comparisons in pairwise testing. For the survival rate, a Pairwise Log-Rank Test was conducted using the R package survival (Therneau 2024).

## Results

### Dual choice experiment

Sucrose vs Sucrose-ToxicantsAnts did not show any preference for pure sucrose (1M) or sucrose with acetone over any of the toxicant-sucrose-acetone mixes at any concentration.

No significant differences were observed between the sucrose solution and any of the toxicant-sucrose solutions (see Table 1 and Figure 2). These results suggest that ants either could not distinguish between the sucrose solution and the toxicant-sucrose solution or had no preference for any one of the solutions.

**Table 1:** Estimated marginal means pairwise comparisons of 1M sucrose/1M sucrose + acetone vs. 1M sucrose + toxicant/1M sucrose + acetone + toxicant.

| Comparison | T-Ratio | P-Value | Sample size (N) |
| --- | --- | --- | --- |
| 1M + acetone 0.12% vs.<br>1M + acetone + 0.12% spinosad 0.015% | 0.830 | 0.409 | 75 |
| 1M + acetone 1.2% vs.<br>1M + acetone 1.2% + spinosad 0.15% | 0.083 | 0.933 | 75 |
| 1M + acetone 0.052% vs.<br>1M + acetone 0.052% + fipronil 0.0001% | 1.038 | 0.301 | 96 |
| 1M + acetone 0.52% vs.<br>1M + acetone 0.52% + fipronil 0.001% | 0.521 | 0.603 | 96 |
| 1M vs.<br>1M + imidacloprid 0.005% | 0.133 | 0.894 | 100 |

**Figure 2:**
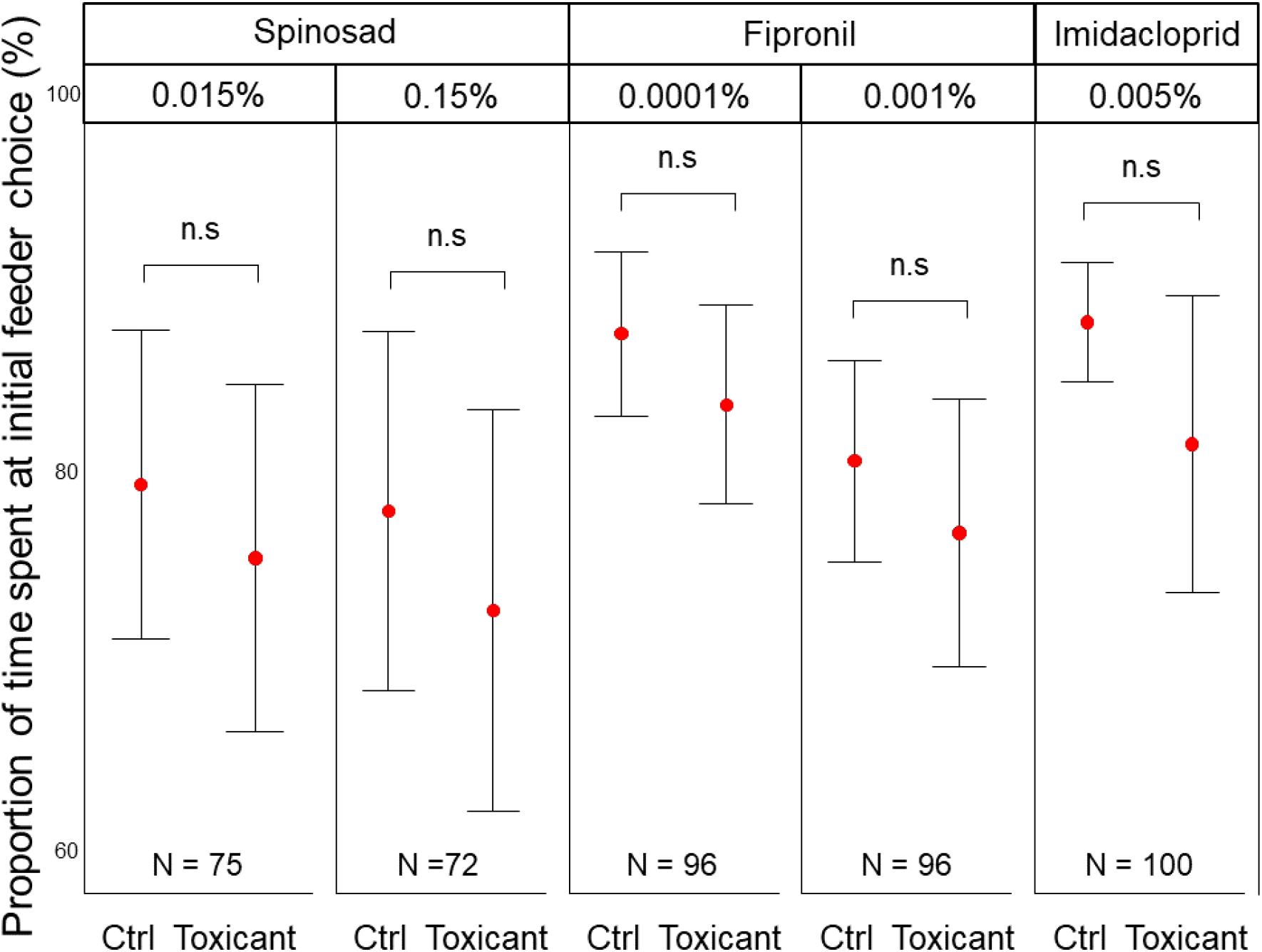
Feeding preference for controls vs. toxicant-laced solutions: Ants showed no preference for pure sucrose/sucrose-acetone solutions (control = Ctrl) over their toxicant or toxicant-acetone counterpart. Error bars represent the 95% confidence intervals. Red dots indicate the means. n.s denote pairs with no statistical differences in preference.

All toxicants at all concentrations resulted in survival after 72 hours of 50% or below (Survived ants per toxicant: spinosad 0.015% and 0.15% = 0%; fipronil 0.0001% and 0.001% = 39% and 3%; imidacloprid 0.005% = 18%, see Figure 3), indicating that the toxicant concentrations used were effective, and comparable with the effectiveness of concentrations used for ant control and eradication attempts in nature. The survival rate after 72 hours for the control group (pure sucrose) was 86%. Pairwise Log-Rank test comparisons showed that all toxicant-sucrose mixes resulted in significantly lower survival rates compared to the control group (Control vs. every single toxicant: p < 0.0001). We found no significant difference in survival between ants fed with a solution containing acetone and those fed with a control solution without acetone over the 72-hour period (GLM: z = 0.023, p = 0.981).

**Figure 3:**
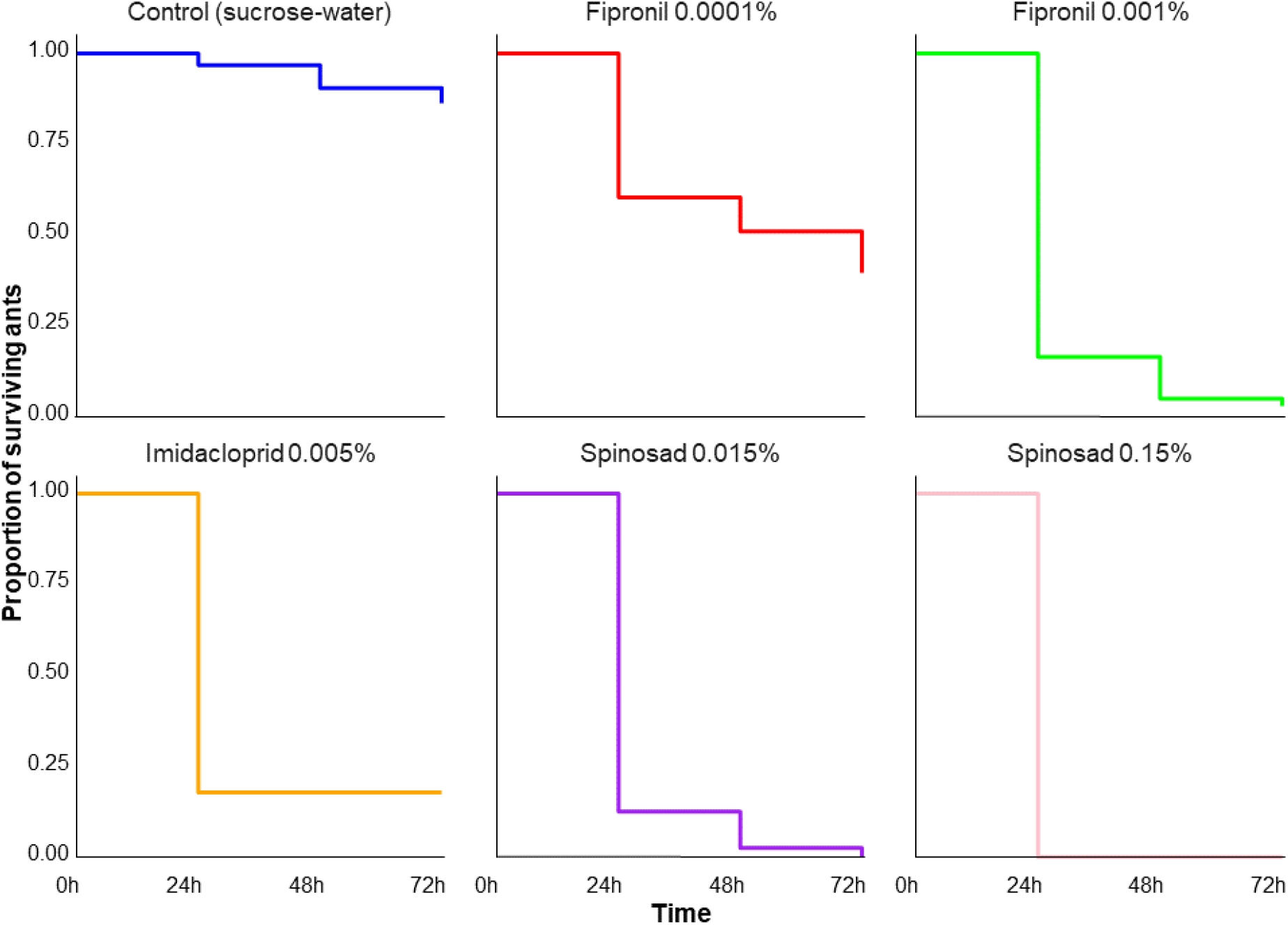
Proportion of surviving ants over time (72h) for different toxicants: All toxicant successful eradiated 50% or more of the tested ants after 72h. The toxicant groups showed a significantly lower survival rate compared to the control group, which was fed with water and sucrose only.

#### Sucrose vs Sucrose-& egg-white protein

Ants showed a strong preference for pure sucrose (1M) over a sucrose-egg-protein mix (25:1 ratio). When pure sucrose was encountered first, ants spent 87% of their total drinking time on it, compared to only 50% when the first choice was the sucrose-egg-protein mix. (t-ratio = 2.710, p = 0.009, see Figure 4).

**Figure 4:**
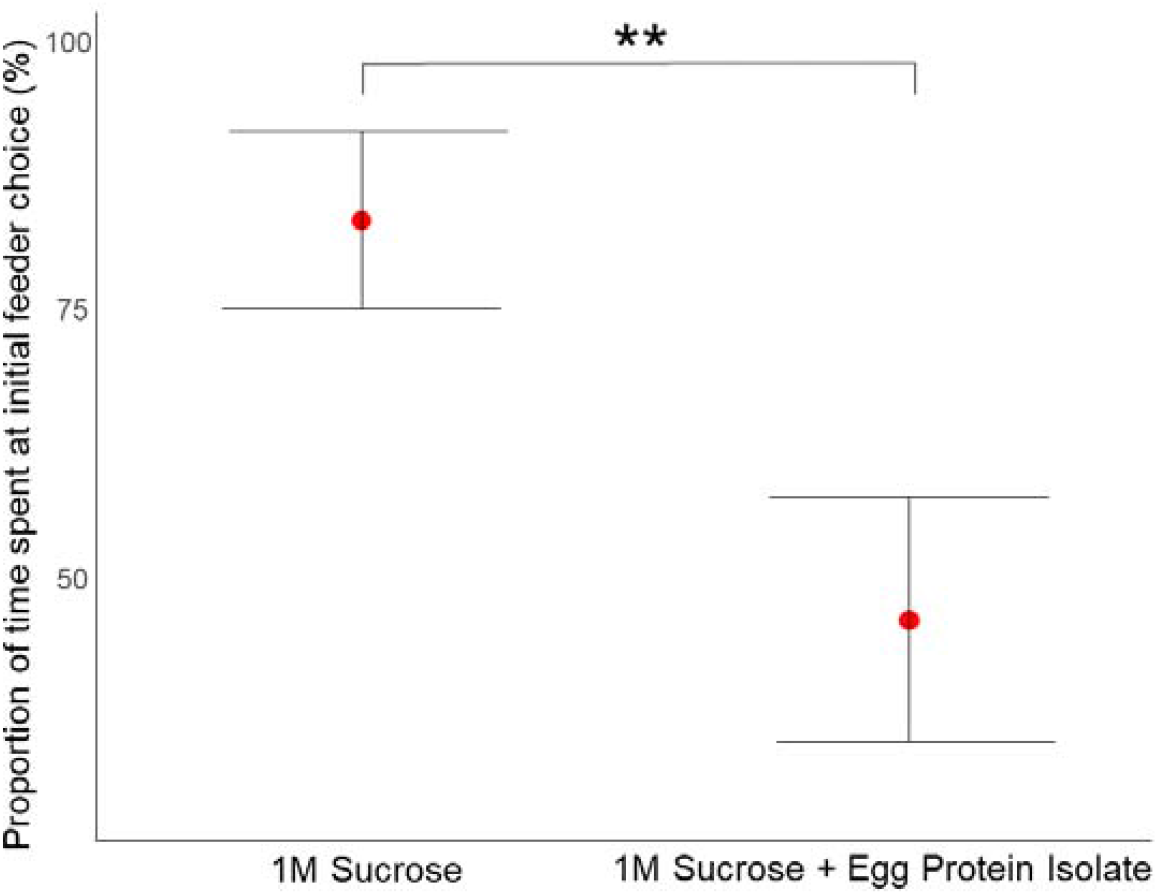
Feeding preference pure sucrose versus a sucrose-protein mix: Ants preferred pure sucrose (1M) over a sucrose + egg-protein mix. Error bars represent the 95% confidence intervals. Red dots indicate the means. Double Asterisks (**) denote a statistically difference between the two solutions.

## Discussion

We demonstrated that *Linepithema humile* did not show any significant difference in their feeding preferences between pure sucrose solutions and sucrose solutions containing one of the three insecticides (fipronil, spinosad, or imidacloprid). Additionally, ants preferred pure sucrose over a sucrose-egg isolate white protein mix.

Since we presented both food sources simultaneously, we can conclude that ants either cannot detect the presence of these toxicants, or if they do, the taste of the toxicant is neutral and does not affect their preference. Given that the comparative nature of the dual feeder method, the lack of a notable preference strongly indicates that the ants were not deterred by the insecticides after one visit. Thus, the toxicant taste itself is likely not the reason why baits containing these toxicants are often rejected over time.

To the best of our knowledge, there are no reports that ants taste toxicants in an otherwise appetitive matrix (e.g., sucrose). It has been shown that hover flies cannot detect neonicotinoids in sucrose solutions by taste (Clem et al. 2020). Additionally, a study on *Apis mellifera* and *Bombus terrestris* showed that their gustatory neurons did not respond to neonicotinoids, including imidacloprid, indicating they cannot taste these chemicals. However, they showed a preference for them, suggesting neonicotinoids have a rewarding effect that does not involve direct tasting (Kessler et al. 2015; Arce et al. 2018). Contrary to these studies, our ants’ lack of preference for our toxicant mixes, especially imidacloprid, showing the absence of a rewarding effect seen in honey bees and bumble bees. One possible explanation is that the sucrose might have had a masking effect on the taste of the toxicants, making them undetectable to the ants when mixed. Alternatively, the toxicants may affect different neurological functions further down the reward and learning pathways, such as by affecting learning, recall, or satiation sensation. It should be noted that Kessler’s two-choice feeding assay ran over 24 hours and in Arce’s assay even 10 days, providing prolonged exposure and potential cumulative effects, while our ants drank only once for a maximum of 90 seconds, which might not be sufficient to detect a long-term reward-enhancing effect. Nevertheless, one common result is that all three studies did not show aversion to the toxicant mix.

We note that our results might depend on the toxicant’s dose. A higher dose might have led to aversion, but as our ultimate aim was to explore the palatability of field-realistic toxicant concentrations, this is not a major concern. Our tests showed that at every dose of every toxicant, ants showed a low survival rate after 72 hours, with rates of 50% or below. Testing the survival rate of ants ensured that they consumed enough toxicant and suffered a survival rate similar to field conditions. We also tested 10 times higher doses for spinosad and fipronil to see if a higher dose might lead to different preferences, but this was not the case. We are thus confident that in field-relevant levels, these toxicants will not be aversive.

One important difference between our laboratory conditions and applied ant control in the field is that ants might receive (in the field) colony feedback influencing their future food source choices (Detrain and Deneubourg 2008; Arenas and Roces 2016a, b). Furthermore, ants can feed multiple times from the baits in nature, unlike in our experiment where they fed only once. In natural settings, ants have to return to the nest after foraging, leading to higher energy consumption and potentially faster toxicant effects (Ma et al. 2024).

We also demonstrated that ants exhibit a clear preference for pure sucrose over a sucrose-egg white isolate mix. This preference could indicate that *L. humile’s* immediate nutritional needs or preferences are better met by pure sucrose, which provides a quick source of energy. Furthermore, it has been shown that a protein-rich diet leads to a decreased worker survival rate (Dussutour and Simpson 2012; Choppin et al. 2023). However, our sucrose-protein mix solution had a high carbohydrate base and is far from being a high protein-rich diet. Additionally, we tested only one protein source (egg-white), which could be a limitation in our study – ants may prefer other types of protein more. Nevertheless, ants may aim to maintain a specific carbohydrate-protein intake ratio as mentioned above. Collecting pure protein or carbohydrates might make it easier for ants to achieve their optimal food intake ratio or to avoid a high protein intake that could lead to reduced worker survival. Nonetheless, our results show that egg-white protein mix is relatively unpalatable to the ants under the conditions tested.

Our finding of egg-white protein being unpalatable in a sucrose mix might not hold true in every situation: If the colony has a protein deficiency, ants might prefer the sucrose-protein mix. Seasonal effects might also play a role; for example, in winter when queens rarely reproduce, ants might prefer pure sucrose, while in summer, during the queen’s main reproductive period, they might prefer the sucrose-protein mix. Studies have shown that ant dietary preferences can vary with seasonal changes and colony needs (Detrain and Deneubourg 2008). Carbohydrate intake can be crucial during certain colony growth phases, while protein is more critical during brood-rearing and reproductive periods (Bazazi et al. 2011, 2016). The protein experiment was conducted between April and June 2023, which is considered a strong reproducing season, hence protein should be needed at this time. This suggests that *L. humile* might prefer pure sucrose over sucrose-protein mixes. We cannot rule out that a bait containing protein, though less attractive than a carbohydrate-only bait formulation, might reach the queen faster due to her protein needs, potentially making the bait more effective.

Given the findings and implications of our study, future research should focus on field studies that mimic natural foraging conditions and account for seasonal variations in dietary preferences. These studies could help refine bait formulations to ensure they are effective across different environmental contexts and stages of colony development. By integrating these findings, pest control strategies can be more precisely tailored to target ant colonies, thereby improving the success rate of ant management programs.

## Acknowledgements

TW was funded by an ERC starting grant to T. Czaczkes, grant number 948181. TJC was funded by a Deutsche Forschungsgemeinschaft Heisenberg Programme grant, grant number 462101190. We would like to thank Silvia Abril for providing ant colonies.

## Data availability statement

The data that supports the findings of this study are available online from https://zenodo.org/records/13482777

## Author contribution

**T. Wagner:** Conceptualization, Methodology, Investigation, Data Collection and Experiments, Writing - Original Draft, Writing - Review & Editing, Supervision, Formal Analysis, Visualisation. **Moana Vorjans:** Data Collection and Experiments, Writing – Review & Editing. **Elias Garsi:** Data Collection and Experiments, Writing – Review & Editing. **Cosmina Werneke:** Data Collection and Experiments, Writing – Review & Editing. **T. J. Czaczkes:** Conceptualization, Methodology, Resources, Visualisation, Writing - Review & Editing, Supervision, Project administration, Funding acquisition.

